# A metabolically engineered bacterium controls autoimmunity by remodeling the pro-inflammatory microenvironment

**DOI:** 10.1101/2022.02.26.482123

**Authors:** Jugal Kishore Das, Fengguang Guo, Carrie Hunt, Shelby Steinmeyer, Julia A Plocica, Koichi S. Kobayashi, Arul Jayaraman, Thomas A Ficht, Robert C. Alaniz, Paul de Figueiredo, Jianxun Song

## Abstract

Immunotherapy has led to impressive advances in the treatment of autoimmune and pro-inflammatory disorders; yet, its clinical outcomes remain limited by a variety of factors including the pro-inflammatory microenvironment (IME). Discovering effective immunomodulatory agents, and the mechanisms by which they control disease, will lead to innovative strategies for enhancing the effectiveness of current immunotherapeutic approaches. We have metabolically engineered an attenuated bacterial strain (i.e., *Brucella melitensis* 16M Δ*vjbR*, BmΔ*vjbR*) to produce indole, a tryptophan metabolite that controls the fate and function of regulatory T cells (T_regs_). We demonstrated that treatment with this strain polarized M2 macrophages (Mφ) which produced anti-inflammatory cytokines (e.g., IL-10) and promoted T_reg_ function; moreover, when combined with adoptive cell transfer (ACT) of T_regs_, a single treatment with our engineered bacterial strain dramatically reduced the incidence and score of autoimmune arthritis and decreased joint damage. These findings show how a metabolically engineered bacterium can constitute a powerful vehicle for improving the efficacy of immunotherapy, defeating autoimmunity and reducing inflammation by remodeling the IME and augmenting T_reg_ function.

## Main

Numerous studies have demonstrated promising results using immunotherapy to treat autoimmune and pro-inflammatory disorders^1-4^. Nevertheless, despite advances in the field of immunotherapy, including regulatory T cell (T_reg_)-based therapy, the efficacy and benefits of these approaches remain less satisfactory due to limited *in vivo* T_reg_ expansion and persistence, the pro-inflammatory microenvironment (IME), and insufficient T_reg_ trafficking to inflamed sites. In addition, immune dysregulation of the IME contributes to disease^5^. Immunotherapeutic combinations may produce greater efficacy, and thus strategies that circumvent these barriers are urgently needed.

To address this need, we developed and tested an intervention that combined two innovations. First, although a therapeutic role for live attenuated bacterial vaccines in addressing infectious diseases is undeniable and appreciated in cancer immunotherapy, the use of bacterial agents to manage autoimmune and pro-inflammatory diseases remains limited. Second, a growing number of microbiota-specific products and metabolites have novel immunologic properties that constitute one mechanism whereby the microbiota influence host health and disease^6,7^. We therefore developed a metabolically engineered bacterial vaccine to produce immunomodulatory metabolites that improve autoimmunity and inflammation.

For the bacterial vector, we selected an attenuated strain of *Brucella melitensis* that harbors a deletion in *vjbR*, a master regulator of virulence (BmΔ*vjbR*)^8^. Like other Gram-negative organisms, *Brucella* strains express a lipopolysaccharide (LPS) lacking endotoxin activity. Importantly, BmΔ*vjbR* has been shown to be safe in immunocompetent and immunocompromised mice^9^, goats^10^, sheep^11^, and non-human primates^12^. We also have shown that BmΔ*vjbR* can combat cancer in a murine model by remodeling the tumor microenvironment (TME) to a pro-inflammatory state^8^. Moreover, when BmΔ*vjbR* treatment was combined with adoptive cell transfer (ACT) of tumor antigen (Ag)-specific CD8^+^ T cells, tumor growth and proliferation were dramatically impaired^8^. Conversely, in the current studies, we engineered BmΔ*vjbR* to express tryptophanase (tnaA); i.e., BmΔ*vjbR::tnaA*, to produce the tryptophan metabolite indole, a molecule that modulates the fate and function of T_regs_^13^.

We have reported that indole, when used at a range of physiologic concentrations, suppresses several inflammatory characteristics in immune and non-immune cells, and also augments T_reg_ differentiation^14^. Consistent with our previous reports, we demonstrated that indole suppressed TNF-α production in CD11b^+^ spleen cells after *E coli* LPS (eLPS) stimulation **(Fig. 1a & Fig. 1b)** and dampened their activation by suppressing Akt and ERK signaling pathways in response to microbial agonists (eLPS and heat-inactivated *Salmonella* Typhimurium [HKST]) **(Fig. S1a)**. In addition, indole augmented the differentiation of naive CD4^+^CD25^-^ T cells into induced T_regs_ (iT_regs_) measured by FoxP3 *in vitro* in a dose dependent manner **(Fig. 1c & Fig. S1f**). These findings were consistent with our earlier reports^11^, and were comparable to results from studies using the microbiota metabolite butyrate, albeit with distinct dose-dependency^15^. Based on these findings, we hypothesized that indole would ameliorate immune-mediated inflammation in autoimmune and pro-inflammatory diseases.

**Fig. 1:**
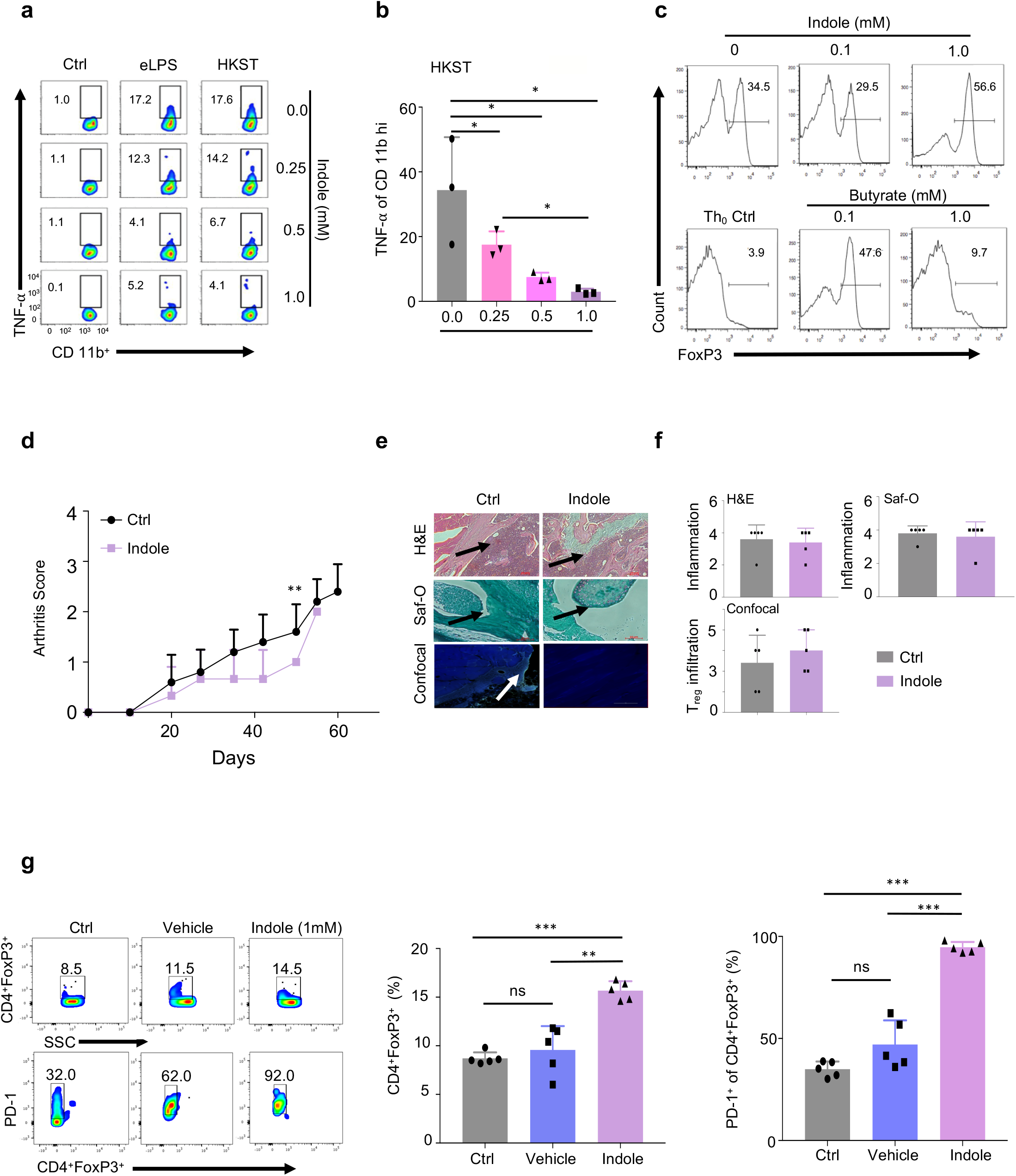
Indole treatment dampens inflammation and promotes T_reg_ expansion and activity. **a**, Indole reduces production of pro-inflammatory TNF-α in CD11b^+^ cells. 0.25, 0.5, or 1.0 mM Indole was dissolved in DMF for the representative experimental flowcytometric analysis. **b**, Graphical representation of flow cytometric dot-plots derived from 3 independent experiments of HKST group. **c**, Indole promotes dose-dependent differentiation of T_regs_; Experiment (N=3) was performed under T_reg_ skew conditions ([TGF-β] = 2 ng/mL, [IL-2] = 100 U/mL). Th_0_ control represents non-T_reg_ skew conditions. Butyrate was used as a control metabolite. **d**, Effects of indole alone on CIA in mice (N=5). **e**, Representative images of H&E, Safranin O (Saf-O) stained tissues, and confocal microscopy of knee tissues of CIA mice on day 60 post induction of arthritis. **f**, Quantitative analysis of H&E, Saf-O and T_reg_ infiltration from confocal microscopy sections of Control (Ctrl) and indole-treated mice. **g**, Flow cytometric dot-plot analysis of PD-1 and FoxP3 in indole-treated, *ex vivo* stimulated CD4^+^ T cells derived from the LNs and spleen and activated with anti-CD3/CD28 Abs. Exposure to indole drives these cells towards higher T_reg_ phenotype by increased FoxP3 expression. Graphical representation of CD4^+^ T cells (%) derived from the flow cytometric dot-plots of CD4^+^ T cells exposed to indole. Graphical representation of CD4^+^ FoxP3^+^ PD-1^+^ T cells (%) from the flow cytometric dot-plots. Data represent means ± SD. Student’s *t*-test or Tukey’s multiple comparisons test was applied for statistical analysis. *, **, ***: significance at p < 0.05, 0.01, 0.001.

Several lines of investigation demonstrated that indole reduces autoimmune responses in a murine collagen-induced arthritis (CIA) model. First, we showed that the severity of CIA was significantly attenuated in indole treated mice, which exhibited clinical scores of 0.8 ± 0.2 (means ± SEM) at (Day 50), compared to 1.6 ± 0.5 in controls (**Fig. 1d)**. However, a single dose of indole only showed a slight decrease in inflammation. Similarly, single dose treatment did not induce significant alterations in the infiltration of T_regs_ as assessed by confocal microscopy **(Fig. 1e & Fig. 1f)**. These findings were in striking contrast to our *ex vivo* experimental findings, which indicated that indole significantly promoted the expansion of CD4^+^FoxP3^+^ T_regs_ and enhanced their activation by increased expression of PD-1 compared to the controls (*p*<0.001), an immunosuppressive molecule, in cells derived from the mouse lymph nodes (LNs) and spleen **(Fig. 1g)**. Based on our *in vivo* and *ex vivo* findings, we hypothesized that the sustained delivery of indole in a bacterial vector may greatly improve the durability of the molecule’s immunomodulatory effects, resulting in an attenuation of autoimmunity and inflammation in CIA. To test this hypothesis, we engineered a safe live-attenuated bacterial strain (BmΔ*vjbR::tnaA*) to constitutively produce indole **(Fig. S1c, Fig. S1d & Fig. S1e**).

First, cytokine array profiling analyses showed that BmΔ*vjbR::tnaA* induced the expression of IL-10 (**Fig. 2a & Fig. S2a)**, which promotes the activities of T_regs_ and reduces autoimmunity and inflammation^16,17^. Strikingly, BmΔ*vjbR::tnaA* also significantly (*p*<0.01) reduced the expression of additional pro-inflammatory cytokines like IL-6, IL-1β and TNF-α in macrophages(Mφ) compared to BmΔ*vjbR* parental strain (**Fig. 2a**). Second, we found that BmΔ*vjbR::tnaA*, when co-cultured with bone marrow-derived Mφ (BMDMs), not only significantly reduced the total CD4^+^ T cells (*p*<0.001) but also reduced the production of the pro-inflammatory cytokines such as TNF-α and IFN-γ (*p*<0.001) compared to the BmΔ*vjbR* parental strain **(Fig. 2b)**. Moreover, BmΔ*vjbR::tnaA* promoted the expansion of T_regs_ and significantly enhanced their activity as assessed by IL-10 production (*p*<0.001) and PD-1 expression (*p*<0.01) **(Fig. S2a**). Third, in the CIA model, a significant reduction in arthritis score and incidence was observed following treatment with BmΔ*vjbR::tnaA*. This amelioration of autoimmunity and inflammation was further augmented when BmΔ*vjbR::tnaA* treatment was combined with ACT of T_regs_ **(Fig. 2c)**. Fourth, we observed significantly (*p*<0.01) reduced numbers of infiltrating inflammatory cells into the joints of mice treated with BmΔ*vjbR::tnaA*. This effect was further enhanced by BmΔ*vjbR::tnaA* treatment followed with ACT of T_regs_ compared to the controls (*p*<0.001) **(Fig. 2d)**. Finally, mice treated with BmΔ*vjbR::tnaA* showed reduced infiltrates in the joint as evidenced by H&E analysis and Safranin O (Saf-O) staining of knee cross-sections (60 days post collagen administration). Notably, these findings were further attenuated by addition of ACT of T_regs_ **(Fig. 2d)**. There was also a significant reduction in the total CD4^+^ T cell proportions and a dramatic increase in T_reg_ (*p*<0.001) proportions in mice treated with BmΔ*vjbR::tnaA* compared to controls **(Fig. 2e**).

**Fig. 2:**
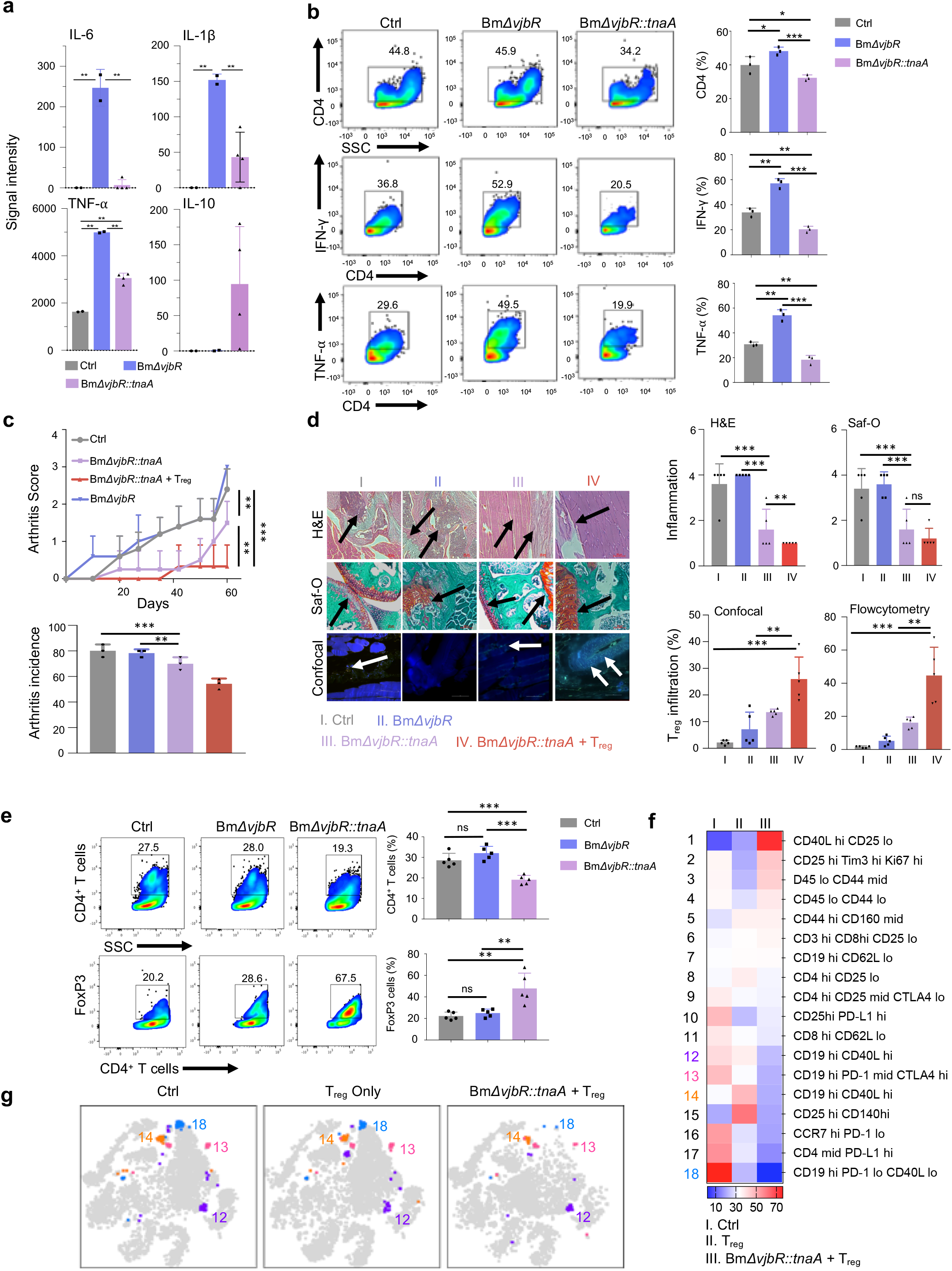
Bm*ΔvjbR::tnaA* significantly dampens inflammation and reduces arthritis in murine CIA model which is augmented by adoptive cell transfer (ACT) of T_regs_. **a**, Cytokine arrays were used to measure pro-inflammatory cytokines produced by control, Bm*ΔvjbR*, and BmΔ*vjbR::tnaA* treated BMDMs. **b**, BMDMs were treated with either BmΔ*vjbR::tnaA* or Bm*ΔvjbR* and then co-cultured with T cells derived from pooled LNs and spleen of C57BL/6 mice. Flow cytometric assays were then performed to measure IFN-γ and TNF-α of T cells. **c**, Arthritis score and arthritis incidence in CIA C57BL/6 mice from control (Ctrl, gray bar); Bm*ΔvjbR* (blue bar); BmΔ*vjbR::tnaA* (pink bar) and BmΔ*vjbR::tnaA* followed by ACT of T_regs_ (T_reg_, red bar) (N=5 in each group). **d**, Representative images of H&E, Saf-O staining, and confocal microscopy from mouse knees on day 60 post CIA induction. Quantitative analysis of T_reg_ infiltration and inflammation scores from these mice are also shown. **e**, Cells from the LNs and spleen were collected from CIA-induced mouse groups (Ctrl, BmΔ*vjbR::tnaA*, and BmΔ*vjbR::tnaA* combined with ACT of T_regs_. These cells were then stained and quantified by flow cytometry using markers for CD4^+^ T cells and intracellular staining of FoxP3 (T_regs_). **f**, CIA-induced mice were treated with PBS (Ctrl), ACT of T_regs_ only (T_regs_ only; N=5), or BmΔ*vjbR::tnaA* combined with ACT of T_regs_ (N=5). Cells from the knee and ankle joints were stained with 21 markers and measured by CyTEK aurora flow cytometry. Heatmap shows immune cell profiles in different treatment groups of mice (scale bar represents percentage of cell in each treatment group within each cell type). **g**, viSNE map shows the four subtypes of B cells differentially expressed in the treated group of mice. Data represent means ± SD. Student’s *t*-test or Tukey’s multiple comparisons test was applied for statistical analysis. *, **, ***: significance at p < 0.05, 0.01, 0.001.

To identify the mechanism by which BmΔ*vjbR::tnaA* might be acting, we conducted a multiparametric CyTEK analysis from the cells isolated from the joints of control, ACT with T_reg_ only, or ACT with T_reg_ plus BmΔ*vjbR::tnaA* groups. We found the BmΔ*vjbR::tnaA* reduced the proportion of B cells **(Fig. 2f, 2g, and Table S1)** in addition to promoting T_reg_ expansion. Overall, our results indicate that BmΔ*vjbR::tnaA* not only remodels the IME and facilitates the expansion and suppressive function of T_regs_ but may also modulate B cell-mediated immunity in the CIA model.

Several strategies have been employed to enhance the efficacy of live attenuated bacterial vaccines. However, bacterial vaccines producing immunomodulatory metabolites that alter the immunological tolerance and the IME have not been previously reported. Our work features both conceptual and methodological innovations. First, this work provides the first description of a live attenuated vaccine whose metabolism has been reprogrammed to amplify anti-autoimmune/inflammation activity. Second, this study provides the first description of how combining a single dose of a live attenuated bacterial vaccination with ACT of T_regs_ can achieve potent therapeutic outcomes. Third, we demonstrate how the genetic tractability of BmΔ*vjbR* can be exploited to engineer vaccines with novel properties. Although in this work, we focused on metabolic reprogramming of the bacterium, we envision that future iterations of the BmΔ*vjbR::tnaA* vaccine will include engineered auto Ags to boost persistent auto Ag-specific T_reg_ responses. Fourth, this work showed that indole did not succeed as as a stand-alone agent. However, this limitation was circumvented by delivery using our engineered bacterial vector. BmΔ*vjbR::tnaA* may offer improved pharmacodynamics for indole by sustained production levels and effects *in vivo* versus indole alone. Morever, we showed that the separate individual beneficial effects of Bm*ΔvjbR* and indole act synergistically *in vivo*. Finally, the natural localization of Bm*ΔvjbR* to leukocytes enabled BmΔ*vjbR::tnaA* to provide improved targeted indole delivery and a higher local effective indole concentration in contrast to the untargeted and transient nature of indole as a single agent. In sum, our work presents an attractive new avenue for the development of anti-autoimmunity/inflammation vaccines.

## Methods

### Immunization with CIA

CIA was induced as described previously with minor modifications^18^. Briefly, male C57BL/6 mice were injected with an emulsion of 100 µl of chick type II collagen (Chondrex; 100 µg) in Complete Freund’s Adjuvant (CFA; Chondrex) using a glass tuberculin syringe with 26-gauge needle. The mice were then assessed for development of joint inflammation and clinical arthritis score until Day 60.

### Bacterial culture

BmΔ*vjbR* or BmΔ*vjbR::tnaA* were cultivated and prepared for experimentation as previously described^8^.

### Engineering indole-producing BmΔ*vjbR::tnaA* strain

To generate an indole producing attenuated BmΔ*vjbR* strain, we cloned an *Escherichia coli* (*E. coli*) *tnaA* gene into a broad range bacteria expression plasmid (pBBR1MCS6Y)^19^ and transferred the plasmid to BmΔ*vjbR*.

### BmΔ*vjbR::tnaA* treatment

CIA was induced in male C57BL/6 mice. On Day 7 after the CIA induction, mice were *i*.*v*. injected with 5.0 × 10^7^ live BmΔ*vjbR::tnaA* or PBS control. In the BmΔ*vjbR::tnaA* + T_reg_ combinatorial treating experiments, mice (N=5) were adoptively transferred with 2.5 × 10^6^ CD4^+^CD25^+^ T_regs_ derived from donor LNs and spleen of naive C57BL/6 mice, one week after the BmΔ*vjbR::tnaA* administration.

### Flow cytometric analysis

Cell staining and flow cytometric analysis were performed as described previously^8^ using the described labeling reagents. Briefly, surface and intracellular staining was performed on the single-cell suspensions and analyzed using LSR Fortessa cell analyzer (BD). The joints were also processed and stained similarly with atibodoies listed in Table S2, and data was acquired on CyTEK aurora flowcytometer (Cytek Biosciences). For multiparametric analyses, the data were analyzed with FlowJo v10 and represented as heatmaps and tSNE plots.

### Histology and immunofluorescence

Mice were humanely sacrificed on day 60 after induction of CIA, and tissue sections were analyzed as previously described^18^. Briefly, the hind foot paws and knees were removed and fixed in 10% formalin and decalcified in Formical-4 (Decal chemical, Tallman, NY). The fixed tissue sections were then stained with H&E and/or Safranin O fast green (Saf-O) stain. The H&E and Saf-O stained sections were then assessed by semiquantitative system of 0 to 4 as described previously.

Immunofluorescent staining and microscopy were performed on the deparaffinized sections by using FITC anti-mouse FoxP3 antibody (Ab) for T_regs_ and DAPI as nuclear stain.

### Statistical Analysis

Student’s *t*-test was performed for statistical analysis between the groups. All analyses were performed in GraphPad Prism v9. A *p* value of < 0.05 was considered statistically significant.

## AUTHORS’ CONTRIBUTIONS

JS, PDF, RCA and TF conceived and designed the experiments. JKD and FG performed the experiments and analyzed the data. JS, PDF, JKD, FG, RCA, TF, CH, SH, JAP, KSK and AJ wrote the manuscript and provided critical feedback. JS and PDF supervised the research. All the authors read and approved the final manuscript.

## ACKNOWLEDGMENTS

We thank Robbie Moore from COM-CAF, Malea Murphy from IMIL, Sankar P. Chaki from TAMH-CVM, and Elizabeth Bustamante from TAMU-HSC, and the Immunomonitoring Core facility at Houston Methodist Hospital-Texas Medical Center for their technical supports. This work was supported by funding from NIH (R01AI121180 and R01CA221867 to J.S., R01AI110642 to R.A., R01HD084339 to T.A.F., and R01AI141607-01A1 to P.D.F.), NSF DBI1532188, and NSF0854684 to P.D.F.

## Supplementary Figures

**Fig. S1:**
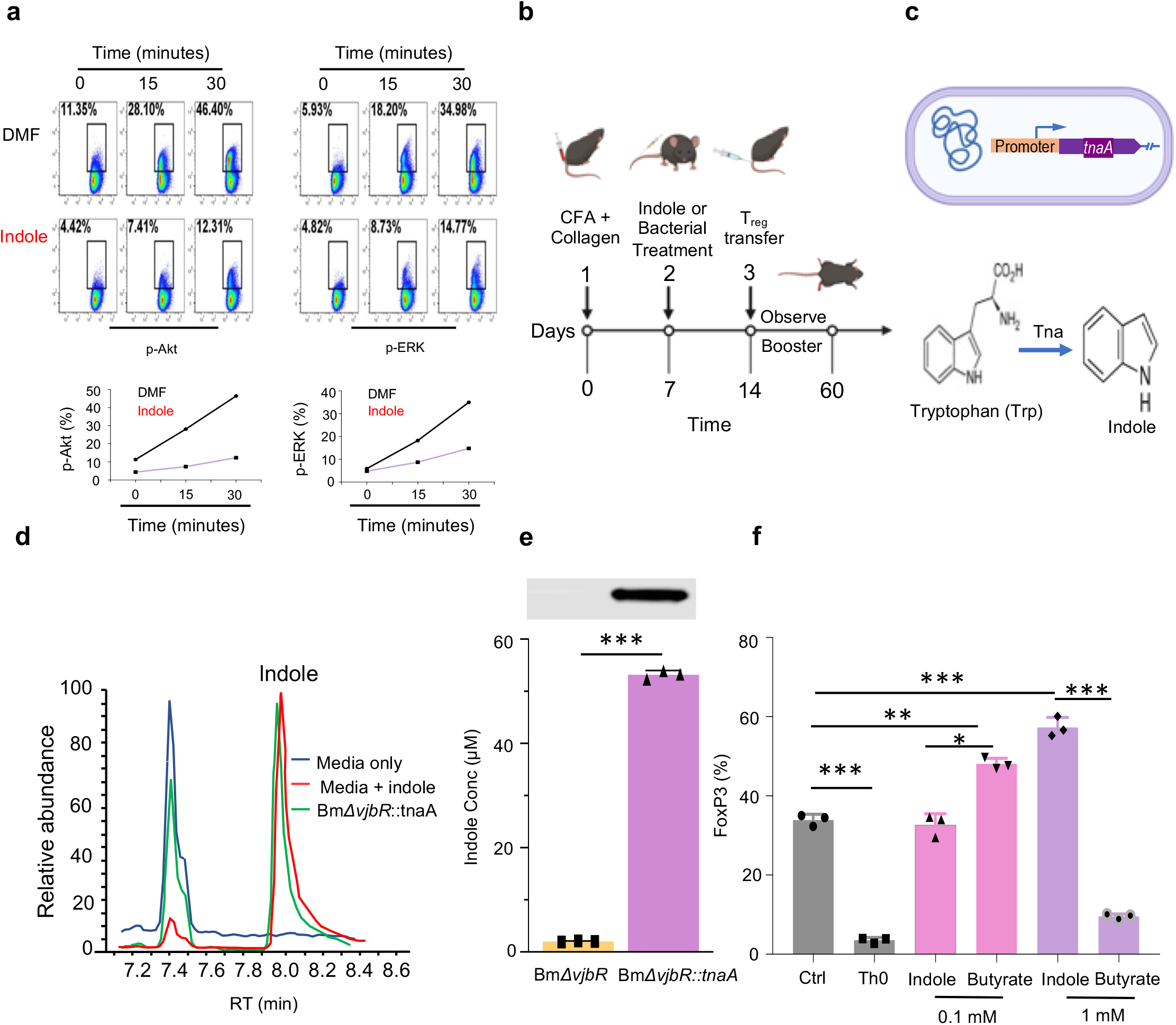
Indole suppresses immune cell activation and Bm*ΔvjBR* is engineered to produce indole. **a**, Indole (1mM) suppresses the Akt and ERK signaling pathways in CD11b^+^ cells in different post incubation time. Flow cytometric dot-plot analysis and graphical representation of Akt and ERK expression with time. **b**, Schematic diagram of murine CIA model used in all the subsequent studies. **c**, Schematic representation of the engineered BmΔ*vjbR::tnaA* harboring a plasmid carrying a *tnaA* expression cassette. The indole biosynthesis pathway is depicted in the figure. TnaA catalyzes the conversion of tryptophan to indole. **d**, Engineered BmΔ*vjbR::tnaA* produced high concentration of indole as shown in the mass-spectrometric analysis. **e**, Western blotting analysis of the expression of tnaA protein in the parental strain compared with the engineered BmΔ*vjbR::tnaA* strain. Graphical representation of the comparative analysis of indole production by BmΔ*vjBR* parental bacterial strain and the engineered BmΔ*vjbR::tnaA* strain. **f**, Indole promotes significantly higher expansion of T_regs_ compared to butyrate at 1mM concentration as shown in the graphical analysis. Graphical data derived from flow cytometric representatives as shown in **Fig. 1C**. Data represent means ± SD. Student’s *t*-test or Tukey’s multiple comparisons test was applied for statistical analysis. *, **, ***: significance at *p*<0.05, 0.01, 0.001.

**Fig. S2:**
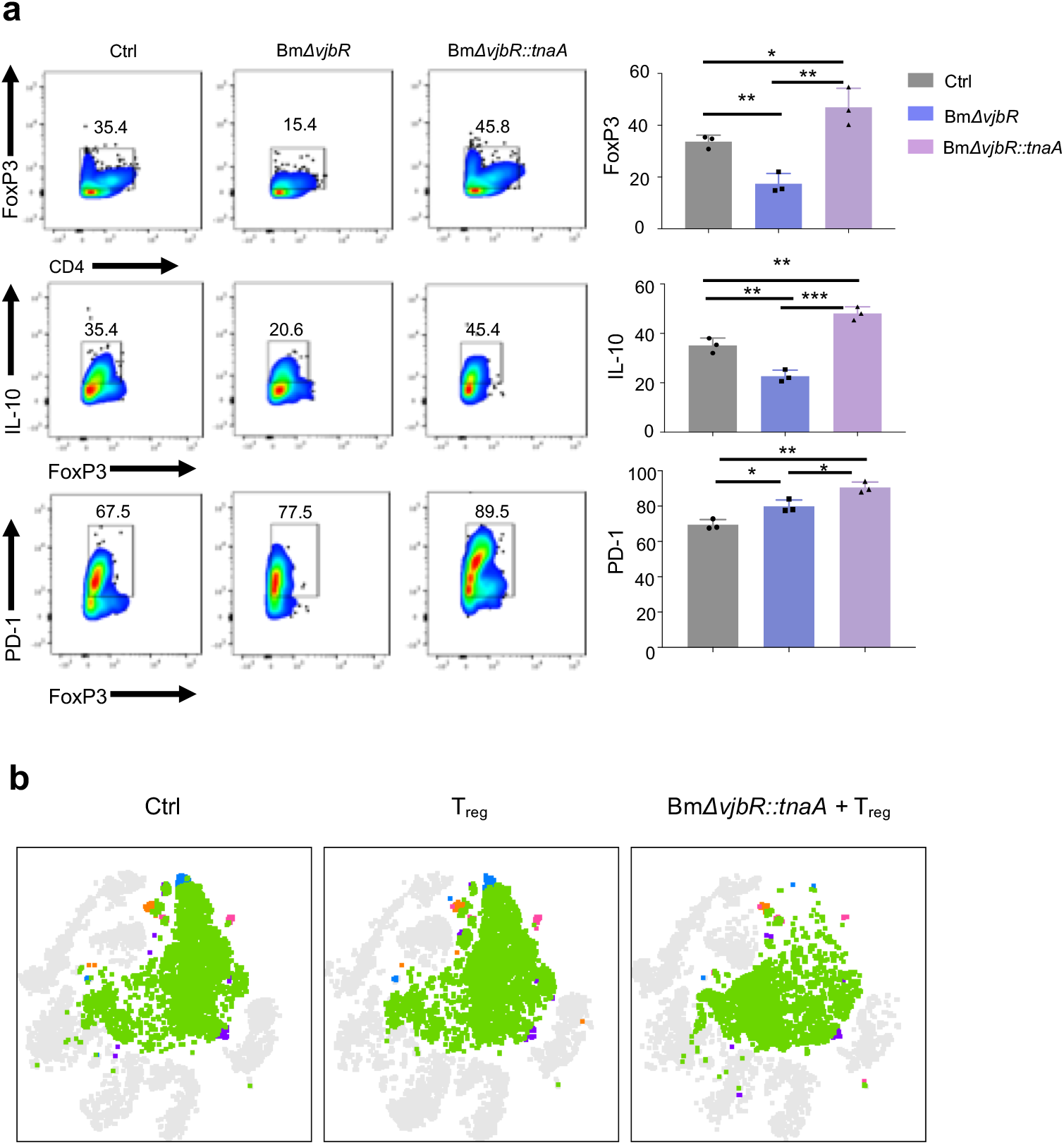
BmΔ*vjbR::tnaA* activates T_regs_ and suppresses B-cell mediated inflammation. **a**, BMDMs infected with either BmΔ*vjbR::tnaA* or Bm*ΔvjbR* were co-cultured with CD4^+^ T cells from mouse LNs and spleen and activated by using anti-CD3/CD28 Abs. Flow cytometric dot-plot assay shows that BmΔ*vjbR::tnaA* treated BMDMs greatly promoted expression of FoxP3 and PD-1 and production of IL-10 in CD4^+^ T cells. The dot-plots are followed by graphical representation from 3 independent experiments. **b**, The total CD19 population is shown in the viSNE plots. Data represent means ± SD. Student’s *t*-test or Tukey’s multiple comparisons test was applied for statistical analysis. *, **, ***: significance at *p*<0.05, 0.01, 0.001.

---

**Supplementary Table 1.**
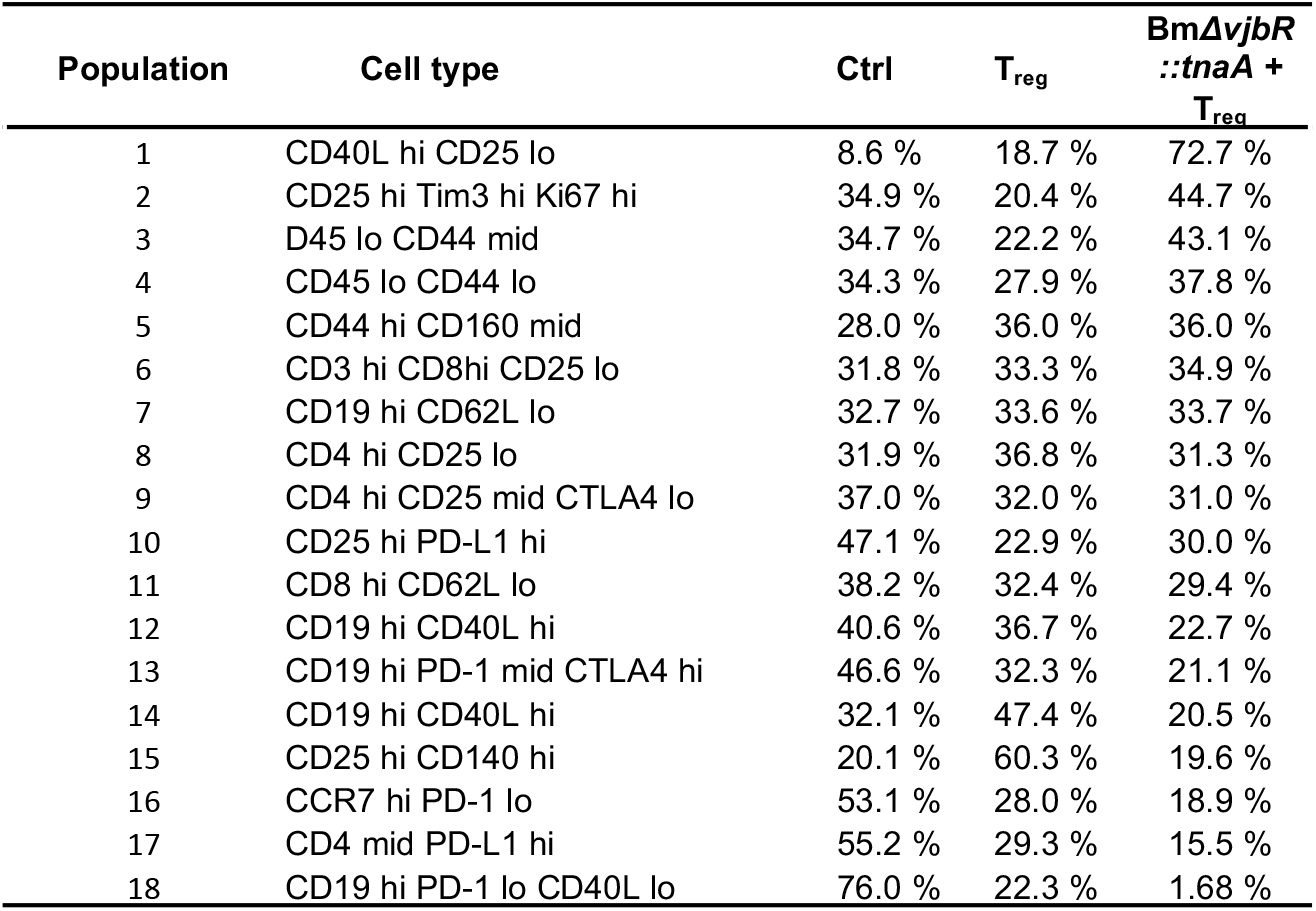
A list of different immune cell types which were differentiated using the tSNE analysis and their proportion in different treatment groups.

| Population | Cell type | Ctrl | T<sub>reg</sub> | BmΔ*vjbR*<br>::*tnaA* +<br>T<sub>reg</sub> |
| --- | --- | --- | --- | --- |
| 1 | CD40L hi CD25 lo | 8.6 % | 18.7 % | 72.7 % |
| 2 | CD25 hi Tim3 hi Ki67 hi | 34.9 % | 20.4 % | 44.7 % |
| 3 | D45 lo CD44 mid | 34.7 % | 22.2 % | 43.1 % |
| 4 | CD45 lo CD44 lo | 34.3 % | 27.9 % | 37.8 % |
| 5 | CD44 hi CD160 mid | 28.0 % | 36.0 % | 36.0 % |
| 6 | CD3 hi CD8hi CD25 lo | 31.8 % | 33.3 % | 34.9 % |
| 7 | CD19 hi CD62L lo | 32.7 % | 33.6 % | 33.7 % |
| 8 | CD4 hi CD25 lo | 31.9 % | 36.8 % | 31.3 % |
| 9 | CD4 hi CD25 mid CTLA4 lo | 37.0 % | 32.0 % | 31.0 % |
| 10 | CD25 hi PD-L1 hi | 47.1 % | 22.9 % | 30.0 % |
| 11 | CD8 hi CD62L lo | 38.2 % | 32.4 % | 29.4 % |
| 12 | CD19 hi CD40L hi | 40.6 % | 36.7 % | 22.7 % |
| 13 | CD19 hi PD-1 mid CTLA4 hi | 46.6 % | 32.3 % | 21.1 % |
| 14 | CD19 hi CD40L hi | 32.1 % | 47.4 % | 20.5 % |
| 15 | CD25 hi CD140 hi | 20.1 % | 60.3 % | 19.6 % |
| 16 | CCR7 hi PD-1 lo | 53.1 % | 28.0 % | 18.9 % |
| 17 | CD4 mid PD-L1 hi | 55.2 % | 29.3 % | 15.5 % |
| 18 | CD19 hi PD-1 lo CD40L lo | 76.0 % | 22.3 % | 1.68 % |

**Supplementary Table 2:** List of Abs used in CyTEK assay

| Ab | Clone | Fluorophore | Source |
| --- | --- | --- | --- |
| Ghost Dye Red 710 |  | Ghost Dye Red 710 | Tonbo Biosciences |
| CD45 | 30-F11 | FITC | Tonbo Biosciences |
| CD8a | 53-6.7 | BUV395 | BD Biosciences |
| CD4 | GK1.5 | eFluor 450 | Thermo Fisher Scientific |
| CD3 | 17A2 | APC-Fire 810 | BioLegend |
| CD19 | 1D3 | BUV805 | BD Biosciences |
| MHC II | M5/114.15.2 | Violet Fluor 500 | Tonbo Biosciences |
| CD160 | 7H1 | APC | BioLegend |
| PD-L1 | MIH5 | Super Bright 780 | Thermo Fisher Scientific |
| CTLA-4 | UC10-4F10-11 | PE-Cy7 | Tonbo Biosciences |
| PD-1 | 29F.1A12 | PE-Dazzle 594 | BioLegend |
| Lag-3 | C9B7W | BUV661 | BD Biosciences |
| TIM-3 | 5D12/TIM-3 | Brilliant Violet 605 | BD Biosciences |
| CD44 | IM7 | PerCP | BioLegend |
| CD62L | HRL1 | BUV737 | BD Biosciences |
| Ki-67 | B56 | Brilliant Violet 711 | BD Biosciences |
| CCR7 | 4B12 | PE | BioLegend |
| CD69 | H1.2F3 | BUV563 | BD Biosciences |
| CD25 | PC61 | Brilliant Violet 650 | BioLegend |
| CD40L | MR1 | BB700 | BD Biosciences |
| CD28 | 37.51 | BUV496 | BD Biosciences |

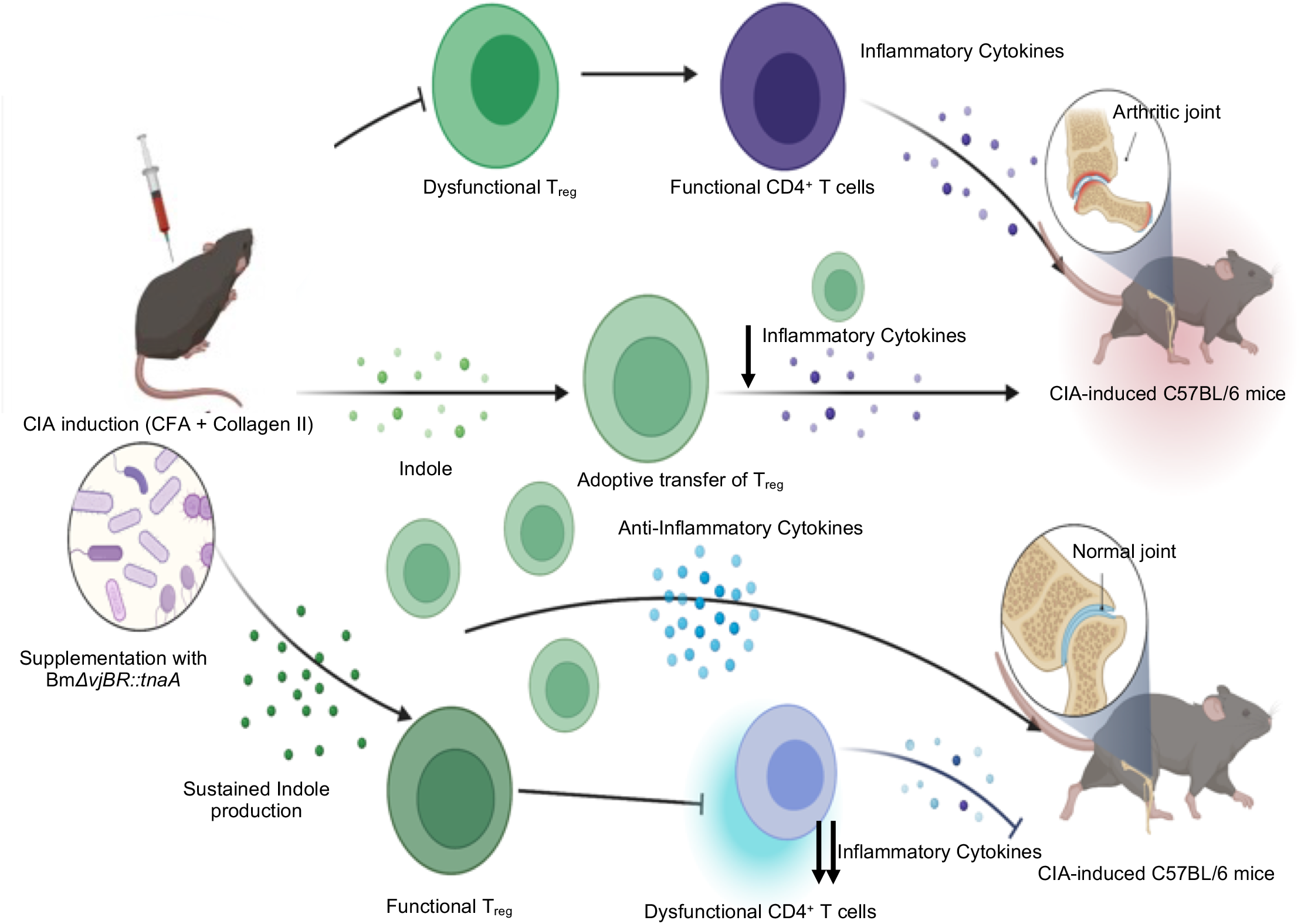

## Schematic representation of amelioration of CIA in C57BL/6 mice

The illustration depicts the process of amelioration of CIA in C57BL/6 mice by the engineered *BmΔvjbR::tnaA* bacterial strain. C57BL/6 mice may harbor a microbiome deficient in production of the metabolite indole. When challenged with the CIA, these mice rapidly develop arthritis due to dysfunctional T_regs_ and rapid expansion of CD4^+^ effector T cells. However, when *BmΔvjbR::tnaA* bacteria are administered to mice bearing CIA the T_reg_ activity is substantially increased and the pathogenic activity of CD4^+^ effector T cells is compromised, which results in amelioration of joint inflammation and improved synonyms. This activity is greatly augmented by adoptive transfer of T_regs_.

